# Methylation-associated mutagenesis underlies variation in the mutation spectrum across eukaryotes

**DOI:** 10.1101/2025.05.28.656604

**Authors:** Fabián Ramos-Almodóvar, Ziyue Gao, Benjamin F. Voight, Iain Mathieson

**Author notes:** Correspondence to BFV or IM. These authors jointly supervised this work (Benjamin F. Voight, Iain Mathieson).

## Abstract

Mutation spectra vary across genetic and environmental contexts, leading to differences between and within species. Most research on mutation spectrum has focused on the trinucleotide (3-mer) mutation types in mammals, limiting the breadth and depth of variation surveyed. In this study, we use whole-genome resequencing data across 108 eukaryotic species – including mammals, fish, plants, and invertebrates – to characterize pentanucleotide (5-mer) non-coding mutation spectra using a Bayesian approach. Our findings reveal cytosine transition mutability at CpG and (among plants) at CHG sites as the main drivers of variation in mutation spectra across eukaryotes, correlating strongly with genomic CpG and CHG depletion. However, despite the influence of methylation on CpG mutability, genome-wide average CpG methylation levels do not predict CpG transition rates across species and CHG methylation does not predict CHG transition rate, indicating unknown genetic or environmental factors influencing mutation rates at methylated cytosines. Together, our results illustrate the pivotal role of mutagenesis in shaping genome composition across eukaryotes and highlight a gap in knowledge about the mechanisms governing mutation rates.

## Introduction

Mutation is a random process, but it is not uniform. DNA sequence context influences where, what type, and how frequently mutations occur^1,2^. One example is the hypermutability of methylated cytosines at CpG dinucleotides in vertebrate genomes, resulting from spontaneous deamination of 5-methylcytosine to thymine^3,4^. Recognizing the importance of sequence context, recent work has focused on inference of context-specific mutation spectra, which quantify relative mutability based on the flanking nucleotides of a mutated base^5–9^. Most studies of mutation spectra have focused on trinucleotide (3-mer) contexts, where the mutation of interest is analyzed along with one adjacent nucleotide on each side. Consideration of the trinucleotide context has facilitated the identification of mutational signatures in cancer genomes as well as characterization of heterogeneity in polymorphism spectrum across human populations^5–7,10–13^. However, extending the analysis to longer sequence contexts (e.g., 5-mer) provides a higher-resolution view and better captures differences in mutation processes^9,14,15^. We recently developed *Baymer*^8^, a Bayesian hierarchical tree approach that facilitates accurate and robust inference of mutation spectra from polymorphism data. The number of events observed in polymorphism data is much larger (∼100M) relative to *de novo* (∼0.1M) datasets^16^, justifying the use of polymorphism data to infer the mutational spectrum, at least in non-coding regions where the effect of selection is expected to be minimal and not context-specific.

Mutation rates differ across species due to genetic and environmental factors, yet the evolutionary forces shaping context-specific mutation rates remain largely unexplored outside of humans, primates and a few vertebrates^6,17–19^. With the expansion of large-scale sequencing initiatives and conservation genomics efforts, population-level polymorphism data are now available for diverse eukaryotic taxa beyond mammals^20–22^. These resources provide an opportunity to investigate mutation spectra across a wider phylogenetic range. For example, while cytosine methylation is almost exclusive to CpG contexts in vertebrates, it is found extensively in non-CpG contexts in plants and other species where we expect to discover distinct mutational spectra. To this end, we leverage whole-genome resequencing data from 108 eukaryotic species, including mammals, birds, fish, plants, and invertebrates, to characterize variation in mutation spectra in extended pentanucleotide (5-mer) sequence context. Using *Baymer*^8^, we infer context-dependent mutation rates from non-coding polymorphisms and assess the biological and evolutionary factors driving mutation spectrum variation.

## Results

### A catalog of polymorphisms in the non-coding genomes of 108 eukaryotic species

The availability of population polymorphism data in diverse species allows us to study the evolution of mutational mechanisms across many previously unexplored lineages^20,21^. We collected publicly available polymorphism data from whole-genome shotgun sequencing for 113 eukaryotic species^23–122^, including 35 mammals, 5 birds, 15 fish, 33 plants, and 12 invertebrates (**Fig. 1A, Supplemental Table 1**). To reduce issues related to genome assembly quality, variant callability, and the effects of selection on coding regions, we masked repetitive elements, low complexity regions, and exons from each species assembly, with the resulting regions subsequently referred to as the accessible non-coding genome. This resulted in polymorphism datasets with 10^5^ – 10^9^ SNPs per species (**Fig. 1B**).

**Fig. 1.**
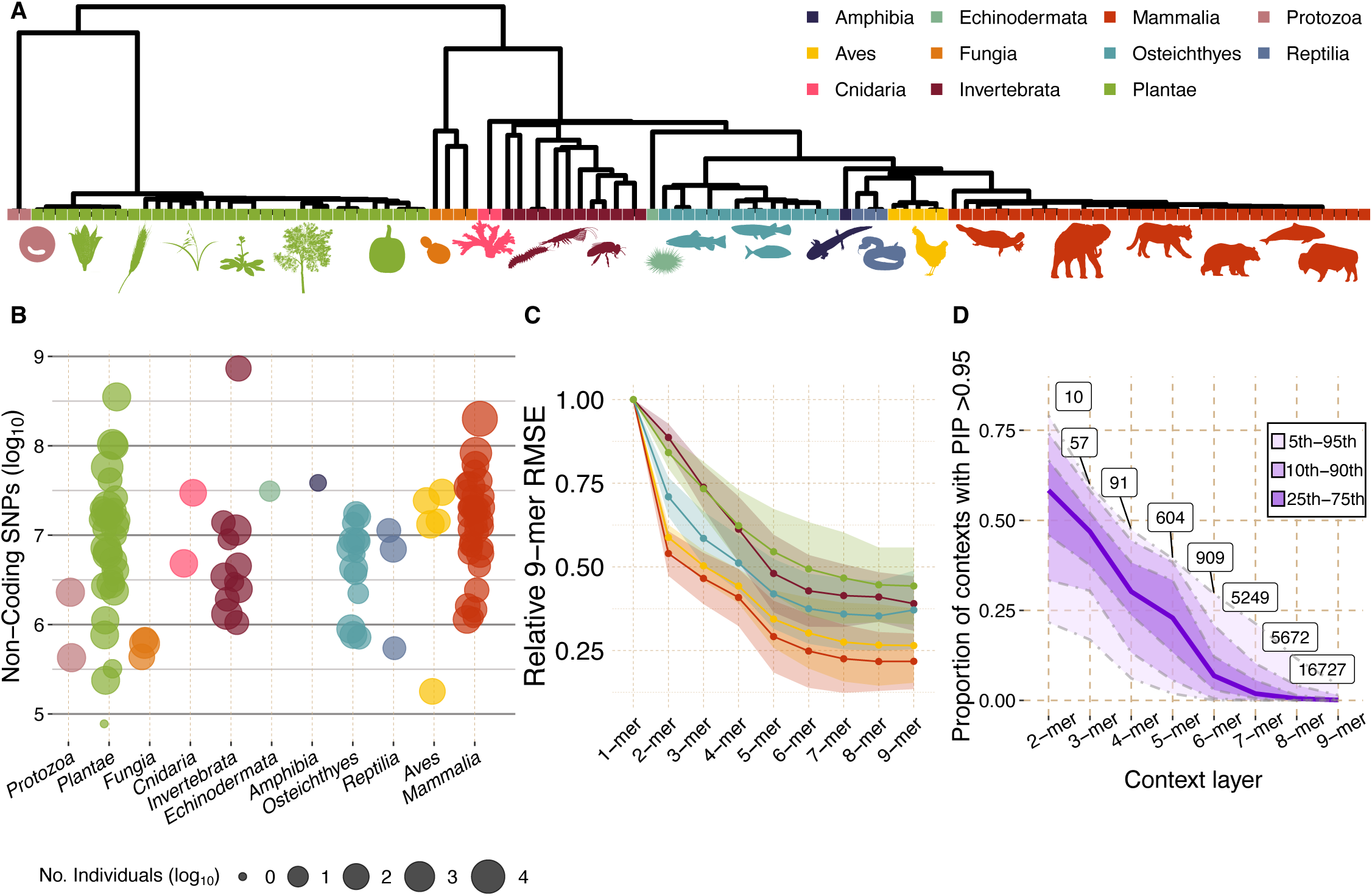
Data summary. **A**) Phylogenetic tree of collected species, generated with TimeTree, with assigned clades labeled at tips. **B**) Filtered single-nucleotide polymorphism (SNP) count (log_10_) per species is shown as individual points, colored by clade and sized by number of individuals (log_10_) in polymorphism dataset. **C**) Root mean square error (RMSE) between a k-mer mutation model trained on odd-numbered basepairs versus a 9-mer mutation model trained on even-numbered basepairs. Points show mean RMSE across species grouped by clade (only clades with n >=5 are shown). Shaded areas show one standard deviation from the mean. **D**) Proportions of contexts with posterior inclusion probability (PIP) greater than 0.95 in each context layer, for every species. Dark line represents the median proportion across species; shaded areas represent labeled quantiles. Number of contexts in 95^th^ percentile is labeled for each layer.

### Baymer captures variation in mutation spectra across species

We used *Baymer*^8^ to infer sequence context windows (up to 4 flanking nucleotides, i.e., ‘9-mers’) separately in each of the 113 species. *Baymer* infers the probability of polymorphism–an approximation to the relative mutation rate of each mutation type–by fitting a hierarchical Bayesian model to the number of observed polymorphisms and the corresponding frequency of the mutable context. A spike-and-slab prior allows contexts to have zero effect on mutation rate, preventing overfitting and allowing the model to adapt to species with very different numbers of observations. We use the default prior and MCMC settings as described in Adams, et al. (2023).

To evaluate the performance of our mutation models within species, we computed the root mean squared error (RMSE) between the mutation models generated from odd- and even-basepairs of the accessible non-coding regions (**Fig. S1**). A higher RMSE suggests higher uncertainty in estimated rates due to technical noise or data sparsity. There is a large drop in RMSE between 1-mer and 2-mer models for mammals and birds due to the inclusion of CpG contexts. For all species, RMSE decreases for larger contexts, but the decrease is substantially slower above 5-mer contexts (**Fig. 1C**). This reflects the fact that the average posterior inclusion probabilities (PIP)–the proportion of contexts with a non-zero effect–drops from 25% at the 5-mer level to 2% at the 7-mer level (**Fig. 1D**). We noted five species with near zero PIPs for every mutation context across every context layer (**Fig. S3**)–a pattern expected in random noise or sparse data that are not informative of mutational processes. We therefore excluded these five species from further analysis and focused on 5-mer contexts for downstream analyses in the 108 remaining species.

### Cytosine transitions at methylation target sites explain most of the variation in mutation spectra

To investigate the variation of context-specific mutation rates in eukaryotes, we first performed principal component analysis (PCA) on the unscaled fitted 5-mer context-specific mutation spectra (**Fig. 2A**), noting that PCA applied to 7-mer (but not 3-mer) contexts produced qualitatively similar results (**Fig. S4**). We found that the first two principal components cluster mutation spectra by clade and capture 84% (PC1: 79.5%, PC2: 4.56%) of the variance in 5-mer mutation spectra across eukaryotes (**Fig. S5A**). The first and second principal components are characterized by variation in NNCGN > NNTGN and NNCHG > NNTHG mutation rates, respectively (where N is any nucleotide and H is any nucleotide other than guanine; **Fig. S5B**). We ruled out saturation of CpG sites due to sequencing of large sample sizes as the source of the captured variation by computing the proportion of CpG sites that are polymorphic for each species (**Fig. S6**).

**Fig. 2.**
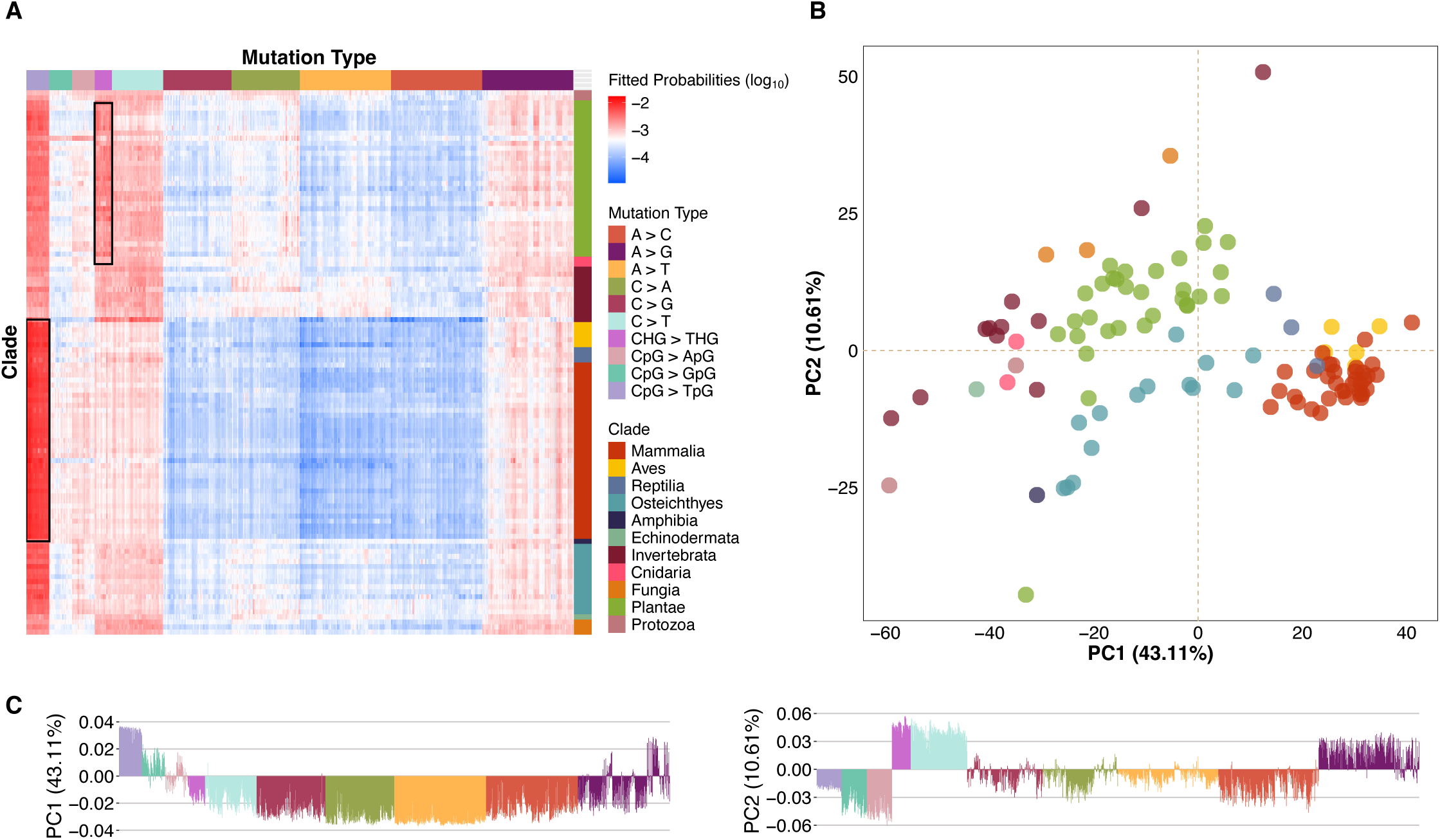
Principal component analysis (PCA) of scaled fitted 5-mer polymorphism probabilities. **A**) Heatmap of raw 5-mer context fitted probabilities; CpG > T and CHG > THG mutations are higlighted in boxes for tetrapods and plants, respectively. **B**) First two principal components from PCA on scaled polymorphism probabilities. Points show each species, colored by clade. **C**) First two principal component loadings, showing individual bars for each mutation type, colored by mutation type.

While the PCA on unscaled mutation spectra reveals variation in absolute contributions to the mutation spectra, it can obscure relative differences due to the disproportionate contribution CpG transitions to the PCA loadings. To assess relative differences in mutation spectra, we performed PCA on scaled 5-mer mutation spectra, where the rates of each mutation type were standardized to have unit variance across species. When examining this relative variation, we found that the first two principal components capture 53.72% (PC1: 43.11%, PC2: 10.61%) of the variance in 5-mer mutation spectra across eukaryotes, and the species still cluster by clade and form a similar phylogenetic cline (**Fig. 2B**). The PC1 of scaled mutation spectra is characterized by variation in NNCGN > NNTGN mutation rate versus other mutation types and PC2 by variation in non-CpG transitions versus other mutation types (**Fig. 2C**). Mammals, birds, and reptiles have the highest weights in PC1, consistent with higher relative CpG mutation rates. The correlation between PCA results and context-specificity of cytosine methylation indicates that most of the variation in context-specific mutation rates across eukaryotes is driven by variation in the rate of cytosine transitions at CpG sites. These results underscore transitions at cytosine methylation target sites as the main driver of context-specific mutation rate variation across eukaryotes.

### CpG transition rates predict genomic CpG depletion but are not predicted by methylation levels

We used the ratio of CpG>T and CpH>T polymorphism probabilities to measure the change in mutation rate due to the CpG context and refer to this as the CpG>T mutation rate ratio. Consistent with the mutagenic effect of methylation, this ratio is largest (5- to 20-fold) in vertebrates, which have high levels of CpG methylation, lower (1.5 to 4.5-fold) in plants and less than 1.5-fold in species without substantial methylation (**Fig. 3A**).

**Fig. 3.**
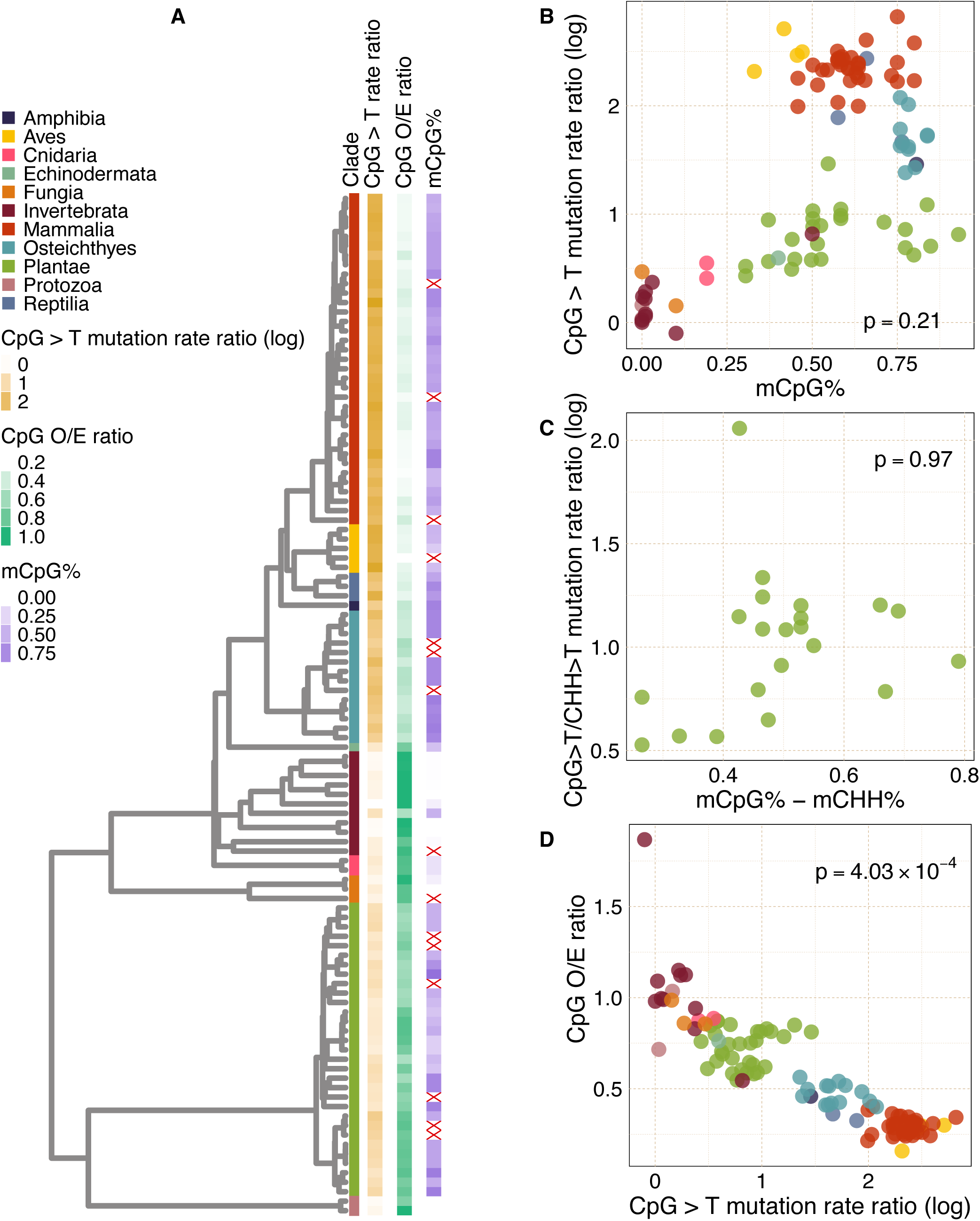
CpG methylation level, mutation rate, and genome composition. **A**) Phylogenetic tree with CpG-related data shown at tips (red crosses denote missing values). **B**) CpG > T mutation rate ratio vs genome-wide average CpG methylation. **C**) Same as B) but restricted to plants and removing the effect of CHG sites on CpG mutation rate ratio and methylation level. **D**) Observed/Expected CpG non-coding genome composition vs CpG > T mutation rate ratio. **B-D**) P-values correspond to phylogenetic regression results.

However, a closer analysis based on genome-wide CpG methylation levels derived from whole-genome bisulfite sequencing^24,123–133^ reveals a more complicated pattern. Phylogenetic linear regression of CpG>T mutation rate ratio on average CpG methylation is not significant (P=0.21; **Fig. 3B**). This lack of relationship is clearly visible among vertebrates, where fish have high methylation levels but low mutation rate ratios, and birds vice versa. We note that the methylation data we collected is derived from somatic tissues. For seven vertebrates where germline methylation data were available^134–139^, we find a Spearman correlation of 0.595 between somatic and germline methylation (**Fig. S7A**) and a relationship that is consistent with the somatic mutation data (i.e. higher methylation but lower CpG mutation in fish relative to mammals; **Fig. S7B**). Thus, presence of CpG methylation entails a higher CpG>T mutation rate ratios, but genome average CpG methylation levels do not predict the magnitude of the ratio. This suggests that some other genetic or environmental factors shape CpG mutation rates across eukaryotes by modifying deamination, repair, or replication-driven error rates at methylated cytosine.

In addition to CpG methylation, many plants have methylation in CHG and CHH contexts. We therefore corrected our analysis of CpG mutability by regressing the CpG>T / CHH>T mutation rate ratio on the difference between CpG and CHH genome-wide average methylation levels in plants (**Fig. 3C**) and observed consistent results (phylogenetic regression P = 0.97).

We next evaluated the effect of the CpG>T mutation rate ratio on CpG content of the accessible non-coding genome, as measured by the ratio of observed CpG fraction among all dinucleotides and the expected CpG fraction based on GC content (CpG O/E ratio). We found that the CpG>T mutation rate ratio strongly predicts the magnitude of CpG depletion (phylogenetic regression P = 4.03x10^−4^) (**Fig. 3D**), with no depletion in species without elevated CpG>T mutation rate ratio; within-clade analysis is reported in **Table S1**. This aligns with the expectation of lower equilibrium genome representation of highly mutable contexts and suggests that CpG content in the non-coding genome is largely determined by the mutation spectrum with minimal influence of selective constraint on CpG content. The one outlier is the honeybee *Apis mellifera*, which has previously been noted to have an exceptionally high genomic CpG content^140^. To provide additional evidence, we examined changes along terminal branches, which offer a cleaner test of this relationship by focusing on recent evolutionary changes. We inferred the CpG O/E and relative mutation rate at the internal nodes of the tree using fastAnc in R and quantified the changes along terminal branches (**Fig S8**). We observed a significant overall negative correlation between shifts in CpG O/E ratio and relative mutation rate, consistent with the expectation that species experiencing recent increases in CpG mutation rates are moving towards lower CpG composition (and vice versa). Although this correlation reaches significance in vertebrates and invertebrates, driven predominantly by honeybee and krill, the negative trend is consistent across other clades (see results in **Table S2**).

### In plants, CHG transition rates predict CHG depletion but are not predicted by methylation levels

To test the relationship between CHG methylation, mutation rate, and genome composition in plants, we extracted genome-wide average CHG methylation levels from whole-genome bisulfite sequencing^123^ and computed the CHG>T / CHH>T mutation rate ratio and the CHG observed-to-expected (O/E) ratio for each species. Analogous to the findings for CpG methylation across eukaryotes, we found that the CHG>T mutation rate ratio is not predicted by CHG methylation (phylogenetic regression P = 0.57, **Fig. 4B**) but does predict the CHG O/E ratio (phylogenetic regression P = 6.75x10^−7^, **Fig. S9**). To investigate whether the same or different factors underlie variation in mutability of CpG and CHG contexts, we regressed our species’ mutation rate ratios on their methylation levels separately for CpG and CHG contexts and found that the residuals were highly correlated (Pearson’s coefficient = 0.829; p = 3.3x10^−6^) (**Fig. 4C**). This suggest that the same factors underlying the unexplained variation in CpG transition rates also underlie variation at CHG sites.

**Fig. 4.**
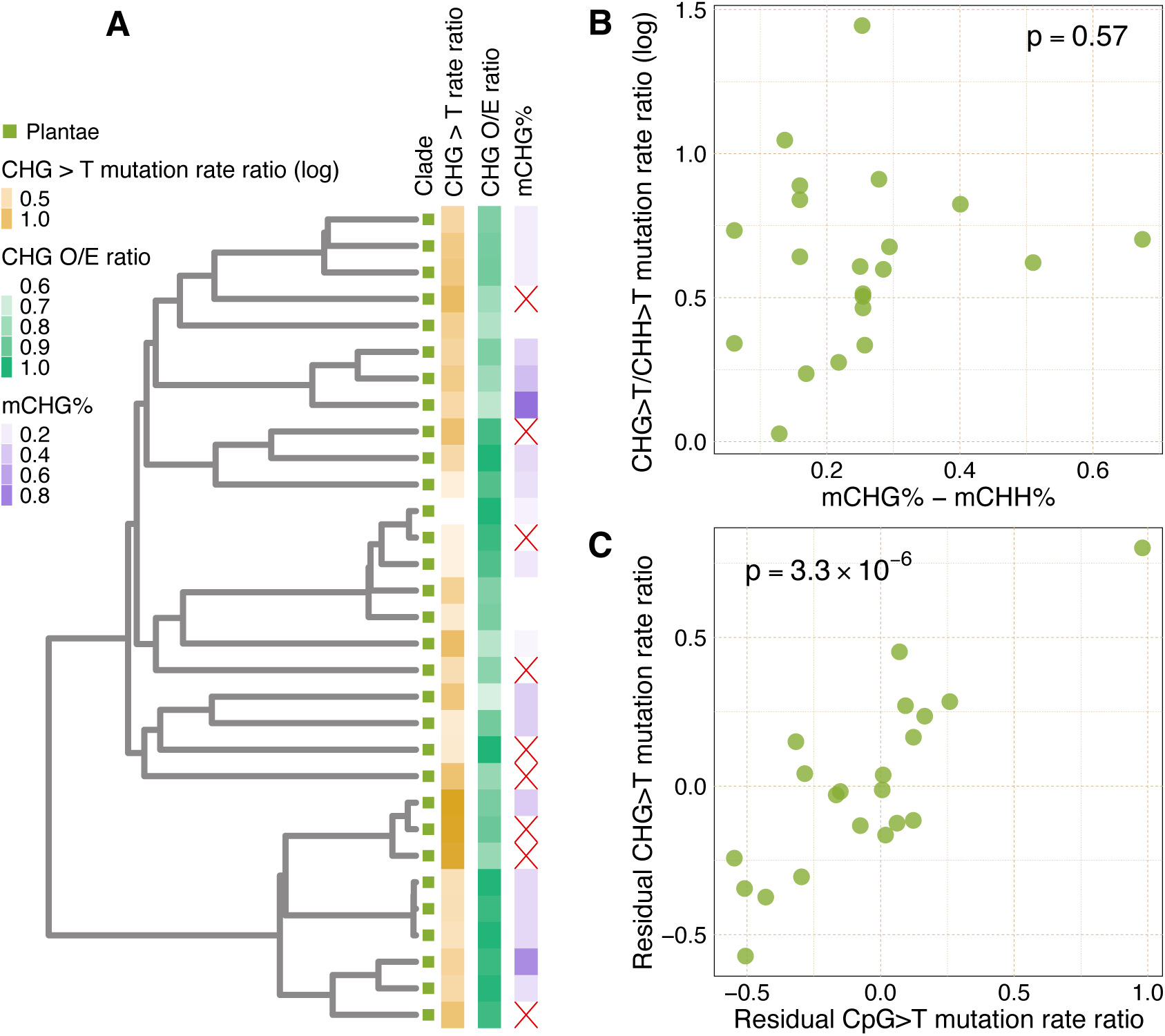
Plant CHG methylation and mutation rate. **A**) Phylogenetic tree with CHG data shown on tips (red crosses denote missing values). **B**) CHG > T mutation rate ratio vs genome-wide average CHG methylation. **C**) Plant CpG and CHG mutation rate ratio residuals from phylogenetic regressions of CpG>T mutation rate ratio on CpG methylation versus those from CHG>T mutation rate ratio regressed on CHG methylation. Plant species are shown as individual points (Pearson’s r = 0.829).

### Mutation signature analysis confirms transitions at methylation target sites as the main driver of mutation spectrum variation in eukaryotes

To corroborate our findings from Baymer-inferred context-specific mutation rates, we used SigFit^141^ to model the observed 5-mer polymorphism spectrum in each species as a linear combination of *k* mutational signatures weighted by differential exposures in each species. Following Beichman et al.^142^, we computed the cosine similarity between the observed and reconstructed mutation spectrum vectors to evaluate model fit to the data. After analyzing the reconstruction performance for *k*=2 to 10, we determined that four mutational signatures had optimal performance and interpretability (**Fig. S10**). These four signatures reflect a CpG transition signature, a CpG and CHG transition signature, and two background signatures (**Fig. 5A**). Consistent with our previous observations, the CpG transition signature was most active in mammals, birds, and reptiles while the signature of CpG and CHG methylation was most active but variable in plants (**Fig. 5B**); the two background signatures are present in all clades.

**Fig. 5.**
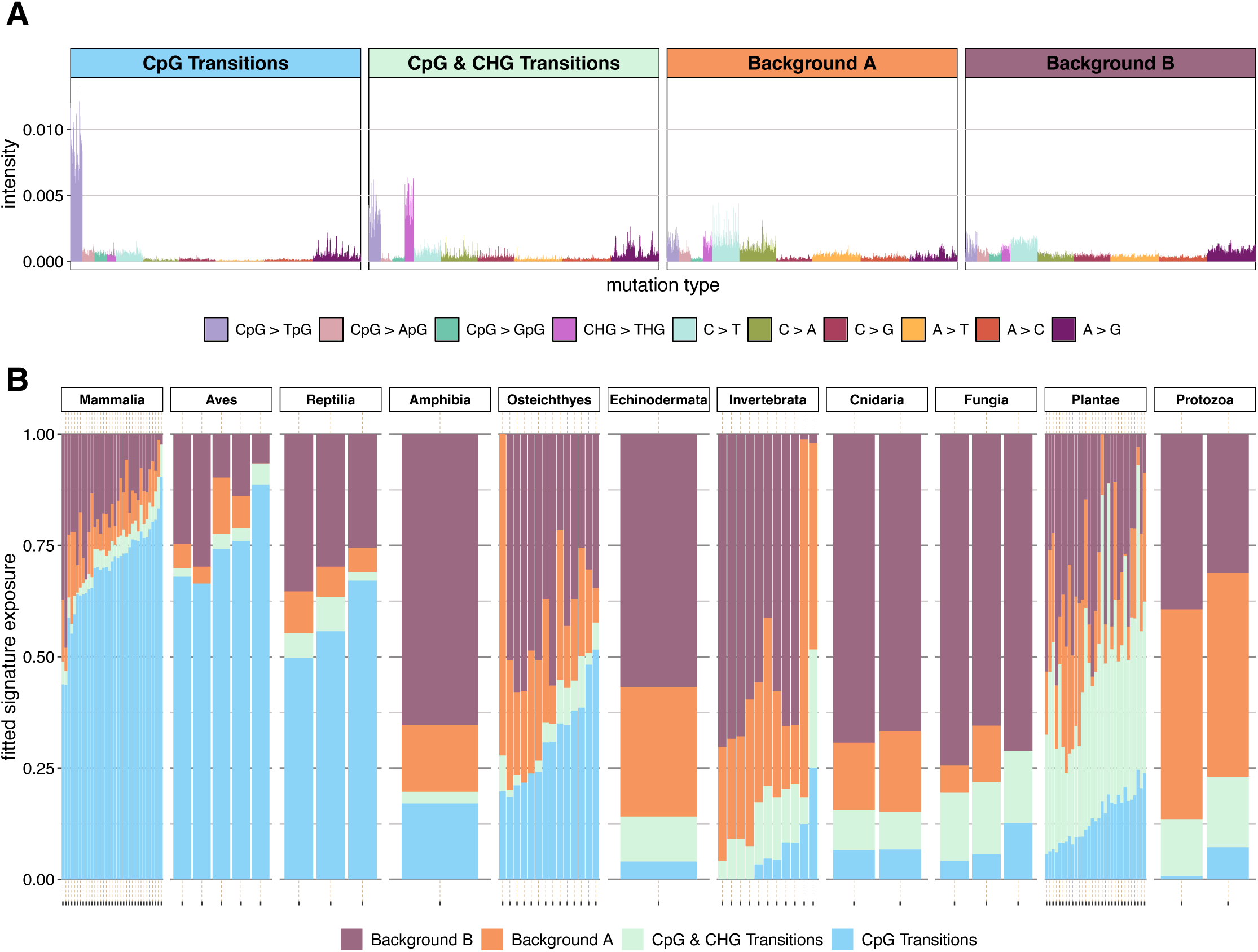
Mutational signature analysis in 5-mer polymorphism spectra. **A**) Extracted signature mutation spectra. **B**) Estimated contributions from each inferred signature to each species’ polymorphism counts.

## Discussion

Our study provides a comprehensive analysis of context-specific mutation spectra across 108 eukaryotic species, leveraging a phylogenetically diverse collection of polymorphism data from whole-genome sequencing. By extending mutation spectrum analysis to 5-mer contexts and looking across a wider range of species, we discover a new axis of mutation spectrum variation, corresponding to transitions at CHG contexts, in addition to the known role of CpG transitions. This is particularly important in plants, where methylation in CHG contexts is common and CHG > T mutations explain a large proportion of mutation spectrum variation.

While our findings reinforce the predominant role of transitions at methylation target sites on mutation spectrum variation across eukaryotic species, they also reveal that methylation is not the only factor at play, as genome-wide average methylation levels do not predict mutation rates at these sites across species after correcting for phylogenetic non-independence. Therefore, while methylation is necessary for high transition rates at CpG and CHG sites, the variation in mutation rates is strongly modulated by other unknown genetic and environmental factors that differ across species and modify rates of deamination, replication error, or repair. One limitation of our current analysis is that it is based on somatic methylation data which might imperfectly reflect methylation patterns in the germline. Consequently, a poor correlation between methylation levels and mutation rates may be observed even if the former strongly determines the latter in the germline. If that is the case, our observations could reflect uncoupling between somatic and germline methylation levels across species. The relationship between mutation and methylation should be reanalyzed when germline methylation data are available for a broader range of eukaryotic species. In contrast, we found that genomic CpG and CHG depletion is highly correlated with cytosine transition rates at CpG and CHG sites, suggesting that these aspects of genomic composition are largely determined by mutational pressures rather than selection, biased gene conversion, or other forces.

In summary, this study advances our understanding of mutation spectrum variation across eukaryotes by integrating 5-mer context mutation spectrum models over a wide range of species. We show that polymorphism data serve as a reliable proxy for mutation data, although absolute mutation rate cannot be assessed and fine-scale variation may be missed due to data artefacts or the lack of polarization of ancestral alleles. Future work should aim to identify genetic or environmental correlates of CpG and CHG transition rates after accounting for methylation, investigate variation across genomic compartments, and sample more broadly from clades with extreme mutational spectra or genome compositions.

## Methods

### Data collection

#### Polymorphism data

We obtained species polymorphism datasets, in the form of .vcf files, from public repositories or directly from authors. We carried out this process by searching through the literature and limiting our selected species to those with polymorphisms called from whole-genome sequencing data, with at least 2x coverage and five individuals (excluding humans, sample size: median=81; mean=259). We also required that the reference genomes used for variant calling had coding region annotations. The full list of species, along with data sources, can be found in **Supplemental Table 1**.

For *Ornithorhynchus anatinus* (Platypus), we aligned 49 individual genomes (in FASTQ format) and called variants using DeepVariant^143^ with its standard protocol (**Supplemental Data**).

#### Reference genome assembly

Corresponding reference assemblies for each species were downloaded from NCBI or publication repository, along with coding and RepeatMasker annotations. **Supplemental Table 1** contains list of sources for polymorphism data and reference genome assembly ID.

### Data preparation

#### Non-coding genome

For each species, we limited the analysis to the accessible non-coding genome by masking out exons, repetitive elements, and low complexity regions. For coding sequences, we extracted genome coordinates (in the form of .bed files) for exons from coding sequence annotation files (downloaded from NCBI or publication repository in the form of .gtf or .gc files) generated from RNA sequencing, for the vast majority of our species (**Supplementary Table 1)**. Repetitive element coordinates were extracted from RepeatMasker annotations. When RepeatMasker annotation was not available, we used *RepeatMasker-4.1.5* with default parameters and specified clade repeat libraries to generate repetitive element annotations. Low-complexity DNA regions were masked using the NCBI tool, *DustMasker-1.0.0* (with -window = 32; default is 64). Our final genomic region for analysis was defined using *bedtools2-2.30.0 getfasta* function by excluding genome coordinates that overlapped exonic, repetitive, and low-complexity coordinates from the above.

#### Polymorphism data

We retained only single-nucleotide polymorphisms (SNPs) from the variant call format (vcf) files with complete 9-mer contexts in the accessible non-coding genome (defined above). Multiallelic sites were treated as independent mutations at the same site. Since the ancestral genome assembly is unavailable for most species, we used reference alleles for polarizing the polymorphisms, assuming the reference allele is the ancestral allele and non-reference allele the mutated allele. Reverse complementary mutation types were combined. To mitigate concerns of high error rate in singletons, we restricted our analysis to SNPs present in at least two individuals, when individual genotypes were available; otherwise, we removed singletons based on non-reference allele count. Because our polymorphism data derive from studies with different sequencing and variant calling strategies, we checked that the PCs of the mutation spectrum were uncorrelated with technical features including SNP count and reference genome N50 (**Fig. S8**).

### Calculation of CpG and CHG observed/expected (O/E) ratio

The observed-over-expected CpG content ratio (CpG_O/E_) was calculated as:

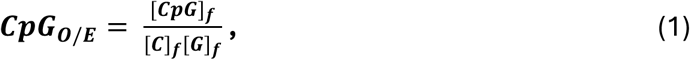

where the observed CpG frequency, [***CpG***]*_f_*, was calculated as the fraction of dinucleotide counts that are CpGs and the expected CpG frequency the product of the observed cytosine and guanine single-nucleotide frequencies, [***C***]*_f_* and [***G***]*_f_*, in the accessible non-coding genome.

The observed-over-expected CHG content ratio (CHG_O/E_) was calculated similarly as:

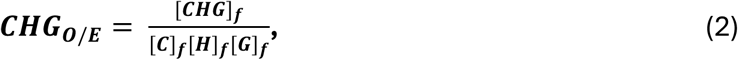

where the observed CHG (where H is any nucleotide other than guanine) frequency, [***CHG***]*_f_* , was calculated as the fraction of trinucleotide counts that are CHGs and the expected frequency the product of the observed frequencies of cytosine, non-guanine, and guanine single-nucleotide, [***C***]*_f_*, [***H***]*_f_*, and [***G***]*_f_*, in the accessible non-coding genome.

### Methylation data collection

#### Cytosine methylation data

We collected averaged genome-wide cytosine methylation levels at CpG sites in each species and at CHG sites in plants from multiple sources (each species’ methylation state and source is indicated in **Supplemental Table 2**) from whole-genome bisulfite sequencing (WGBS) studies^123,124^. Additionally, we collected averaged genome-wide cytosine methylation levels at CpG sites from testes WGBS for seven vertebrate species^134–139^ (listed in **Supplemental Table 2**).

### Context-specific mutation probability models using Baymer

To construct our mutation probability models, we used Baymer^8^, a Bayesian hierarchical tree model approach that estimates context-specific mutation probabilities for increasing context layers, iteratively. Baymer generates regularized mutation probability estimates and provides a measure of uncertainty for each multiplicative shift in mutation probability leading up to the final k-mer mutation context. The context-specific mutation rates for any given context window are calculated as the product of the multiplicative shifts leading to the final mutation context. To generate context-specific mutation probability models from polymorphism data, Baymer requires a table with DNA context counts and polymorphic sites for the largest context size desired for the models (in our case, 9-mers). For each species, we first generated a table of 9-mer context counts within the accessible non-coding genome (defined above). Then, we identified and counted polymorphic sites with complete 9-mer contexts in the same regions. Lower context size counts (e.g. 5-mers) were derived from the 9-mer context size counts.

### Assessing robustness of the mutation model

We applied Baymer to these inputs to generate three models, each using one of three different partitions of the non-coding genome, which we refer to as ALL, ODD, and EVEN. The ALL model consisted of every context and polymorphism in the accessible non-coding genome, while the ODD and EVEN models were only generated using odd or even genome coordinates in the accessible non-coding genome, respectively. The ODD and EVEN models were generated for cross-validation of model performance in each species. We assessed within-species model robustness by computing the root mean square error (RMSE) between the ODD and EVEN polymorphism probability estimates; calculated as:

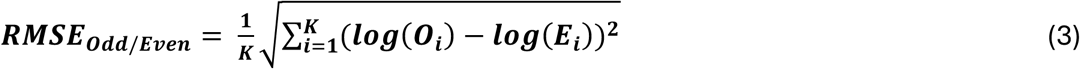

Where **O** and **E** are the fitted context-specific rate vectors, of length K, from the ODD and EVEN models, respectively. **Fig. S1** shows the RMSE_Odd/Even_ for the 5-mer mutation models. Fig. 1C shows the RMSE between 9-mer EVEN mutation models versus k-mer (k = 1 to 9) ODD mutation models, normalized by the k=1 RMSE.

### Mutation rate ratios

Our mutation models infer content-specific relative mutation rates rather than absolute mutation rates. For this reason, we base our mutability analyses on relative differences in context-specific mutation rates, which we refer to as ‘mutation rate ratios’. We estimated these ratios using 5-mer context-specific mutation rates.

CpG>T mutation rate ratio was defined for each species as the ratio of the average C>T mutation rate in all 5-mer CpG contexts over that of CpH contexts (H = A,C,T):

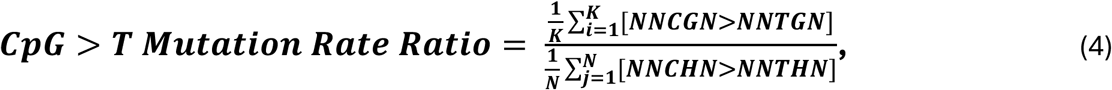

where the summation is across all 5-mer contexts matching the specified patterns.

We note that plant species also experience extensive cytosine methylation at non-CpG contexts. Because of this, the CpG>T mutation rate ratio above may be a biased representation of the relative context-specific CpG>T mutation rate in these plants. To account for this, we defined the CpG>T / CHH>T mutation rate ratio for each plant species as the ratio of the average C>T mutation rate in CpG contexts over that of CHH contexts, removing impacts of methylation in CHG contexts:

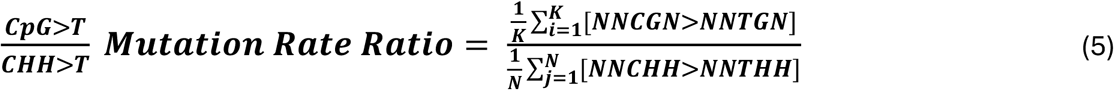

To analyze the relationship between CHG transition rates, methylation, and genome composition in plants, we defined the CHG>T / CHH>T mutation rate ratio for each plant species as the ratio of the average C>T mutation rate in 5-mer CHG contexts over that of CHH contexts:

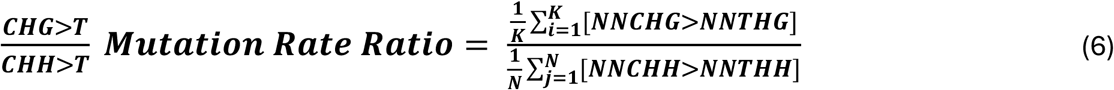

### Phylogenetic tree

Our phylogenetic tree (in Newick file format) was constructed using estimated divergence times from TimeTree (timetree.org). **Supplemental Table 1** contains the list of species names used for TimeTree.

### Principal component analysis (PCA) on raw and scaled fitted mutation probabilities

We performed principal component analysis (PCA) on the raw 3-mer, 5-mer, and 7-mer context size fitted mutation probability models, separately. We scaled the fitted mutation probabilities for each species so that they added up to one and used the prcomp(scale = FALSE, center = TRUE) R (v4.2) function to perform PCA based on single-value decomposition for supplementary figures. For the main text in **Fig. 2**, we performed PCA on scaled 5-mer context mutation probabilities, by standardizing each mutation type to have mean = 0 and standard deviation = 1 across species before PCA; prcomp(scale = TRUE, center = TRUE).

### Phylogenetic linear regressions

To perform analysis accounting for the phylogenic tree structure given the species included in our study, we used the *phylolm-2.6.2* R package phylolm()^144^ function with Pagel’s lambda as a phylogenetic scaling factor and 1000 bootstraps for our regression models. The phylolm() function is an implementation of the phylogenetic generalized least squares regression model:

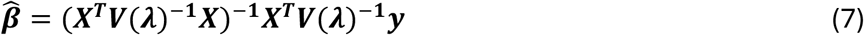

where for *n* species and *k* independent variables, **y** (*n* **×** 1) is the continuous outcome vector, 6 (*n* **×** 1) is the vector of coefficients for the variable matrix **X** (*n* **×** *k*), and **V** (*n* **×** *n*) is the variance-covariance matrix based on the phylogenetic relatedness between the species. Pagel’s lambda is then used to transform **V** into a new matrix, **V(λ)** (*n* **×** *n*), by scaling the oc-diagonal elements of **V** using the scalar **λ**, which is an estimate of phylogenetic signal for the regressed trait. **λ** ranges between 0 and 1, where 1 indicates that the covariance structure is exactly given by the phylogenetic relatedness structure, and 0 indicates no phylogenetic signal for the regressed trait. We report two-tailed P-values.

### Mutational signature analysis with SigFit

We extracted mutational signatures from the observed 5-mer polymorphism spectra using a non-negative matrix factorization approach, implemented in the *SigFit-2.2*^141^ R package. SigFit uses a Bayesian multinomial model to fit specified and/or inferred mutational signatures to observed mutation spectra, an approach equivalent to non-negative matrix factorization. For **G** genomes and **M** mutation contexts, SigFit takes as input a mutation counts matrix (**G** by **M**) and a mutation opportunities matrix (**G** by **M**); the mutation opportunities matrix contains the number of mutable contexts for each given 5-mer mutation type in the species’ genome. For our SigFit analysis, we used similar mutation and context counts as inputs for Baymer inference. We used the extract_signatures() function from SigFit, with iter = 30000 and nsignatures = k (for k = 2,3,…,10). For each SigFit model, we reconstructed the mutation spectra using the fitted mutational signatures and their estimated activity values in each species. We then computed the cosine similarity between the observed and reconstructed 5-mer mutation spectra vectors to measure each model’s performance in each species.

## Data and code availability

Code to reproduce the analysis in this paper is available at https://github.com/framos99/MutationSpectraEvolutionAcrossEukaryotes-Project. Fitted mutation spectra for each species are available at https://doi.org/10.5281/zenodo.15464759. Original sources for data are given in Supplementary Table 1.

## Acknowledgements

F.RA. is grateful for support of the work from the University of Pennsylvania’s Presidential PhD fellowship. This work was supported by NIGMS R35GM133708 (I.M.), R35GM146810 (Z.G.), NIEHS P30ES013508 (B.F.V.), and a Research Fellowship (FG-2021-15702) from the Alfred P. Sloan Foundation (Z.G). The content is solely the responsibility of the authors and does not necessarily represent the official views of the National Institutes of Health.

