## Supplemental Figures for "Methylation-associated mutagenesis underlies variation in the mutation spectrum across eukaryotes"

### Supplementary Figures

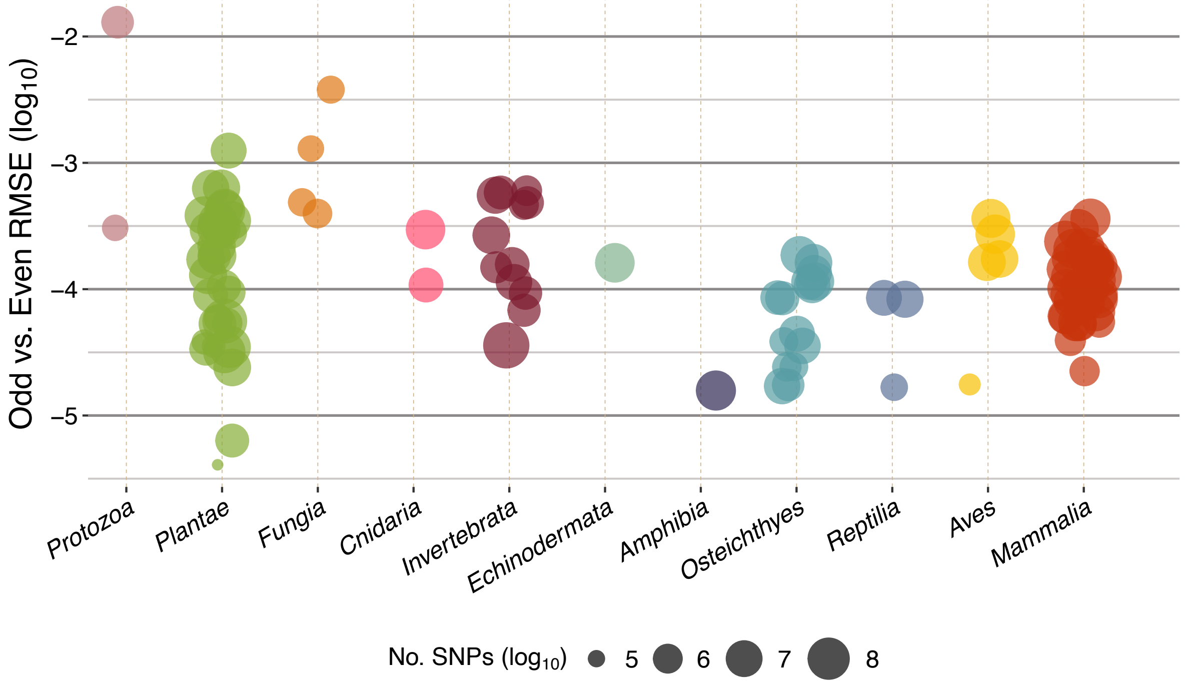


**Fig. S1** | Root mean squared error (RMSE) for within-species Odd versus Even non-coding genome mutation models. RMSE is shown for each species (individual points), colored by clade and sized by single-nucleotide polymorphism (SNP) count ($log_{10}$) in filtered polymorphism dataset.

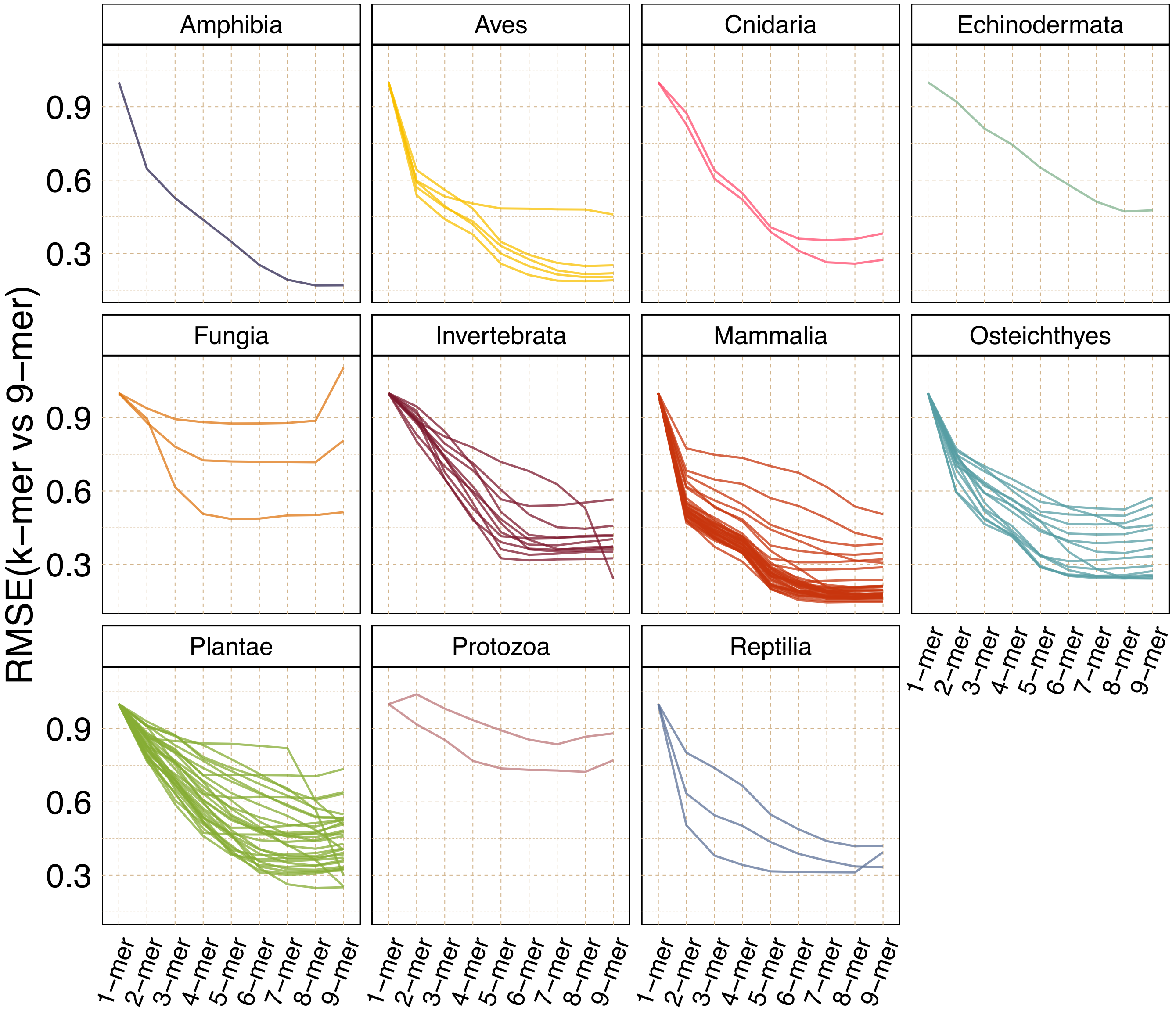


**Fig. S2** | Root mean square error (RMSE) . Each panel shows the proportion of contexts with PIP > 0.95 (on the y-axis) across each context layer size (shown on the x -axis from 2-mer to 9-mer). Species are labeled at the top of the panel and the line color denotes the assigned clade. Five species had near zero proportion of contexts with PIP > 0.95: *Trichaptum abietinum* (Fungia), *Clupea harengus* (Osteichthyes), *Eurytemora affinis* (Invertebrata), *Vitis vinifera* (Plantae), and *Camellia sinensis* (Plantae); these species were not included in our analyses.

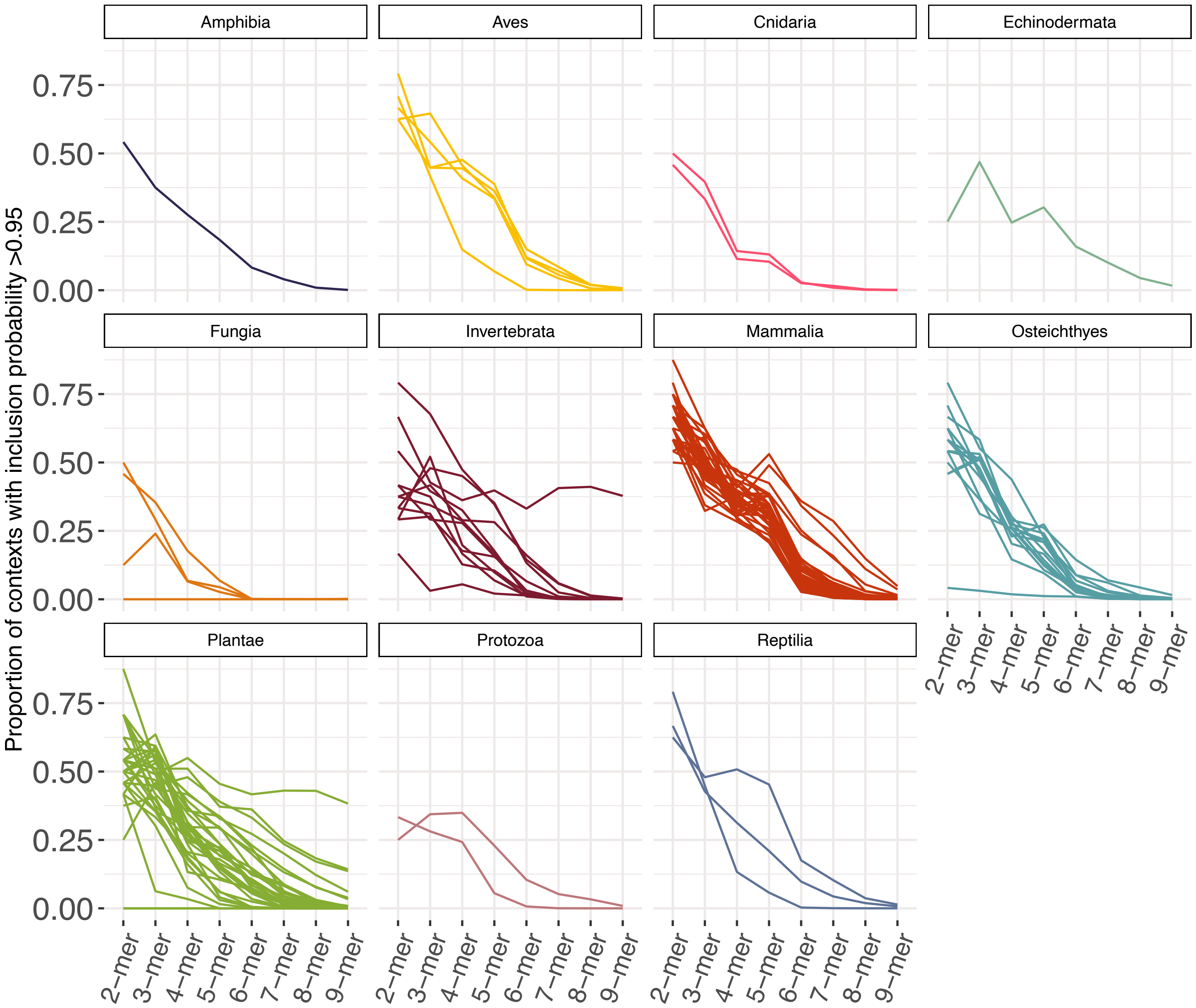


**Fig. S3** | Proportion of contexts with posterior inclusion probabilities (PIP) greater than 0.95 across layers for each species. Each panel shows the proportion of contexts with PIP > 0.95 (on the y-axis) across each context layer size (shown on the x -axis from 2-mer to 9-mer). Species are labeled at the top of the panel and the line color denotes the assigned clade. Five species had near zero proportion of contexts with PIP > 0.95: *Trichaptum abietinum* (Fungia), *Clupea harengus* (Osteichthyes), *Eurytemora affinis* (Invertebrata), *Vitis vinifera* (Plantae), and *Camellia sinensis* (Plantae); these species were not included in our analyses.

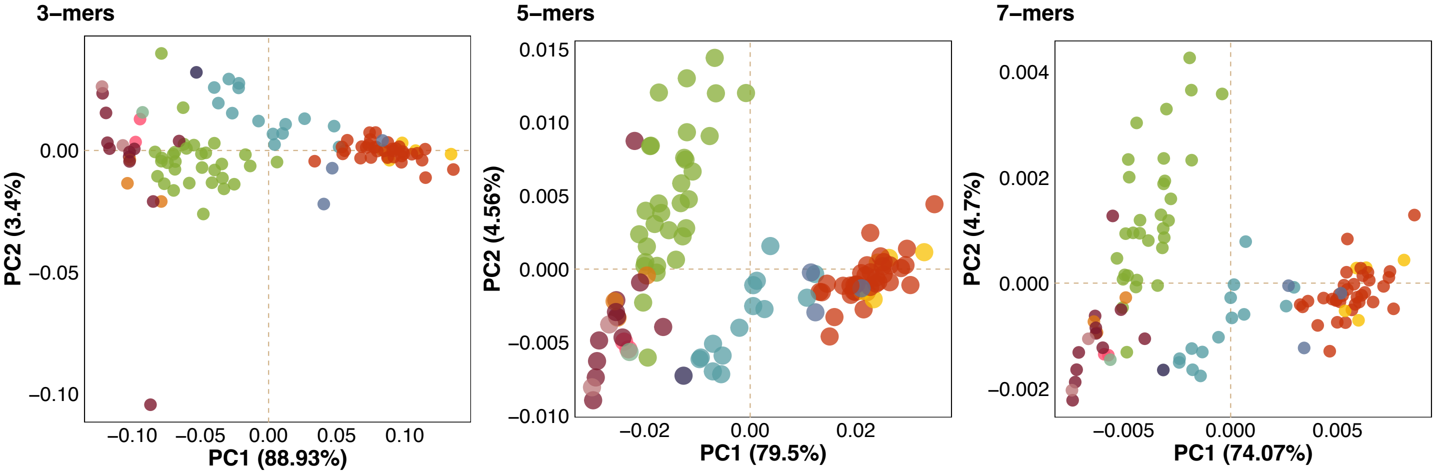


**Fig. S4** | Principal component analysis (PCA) of 3-mer, 5-mer, and 7-mer context fitted polymorphism probabilities.

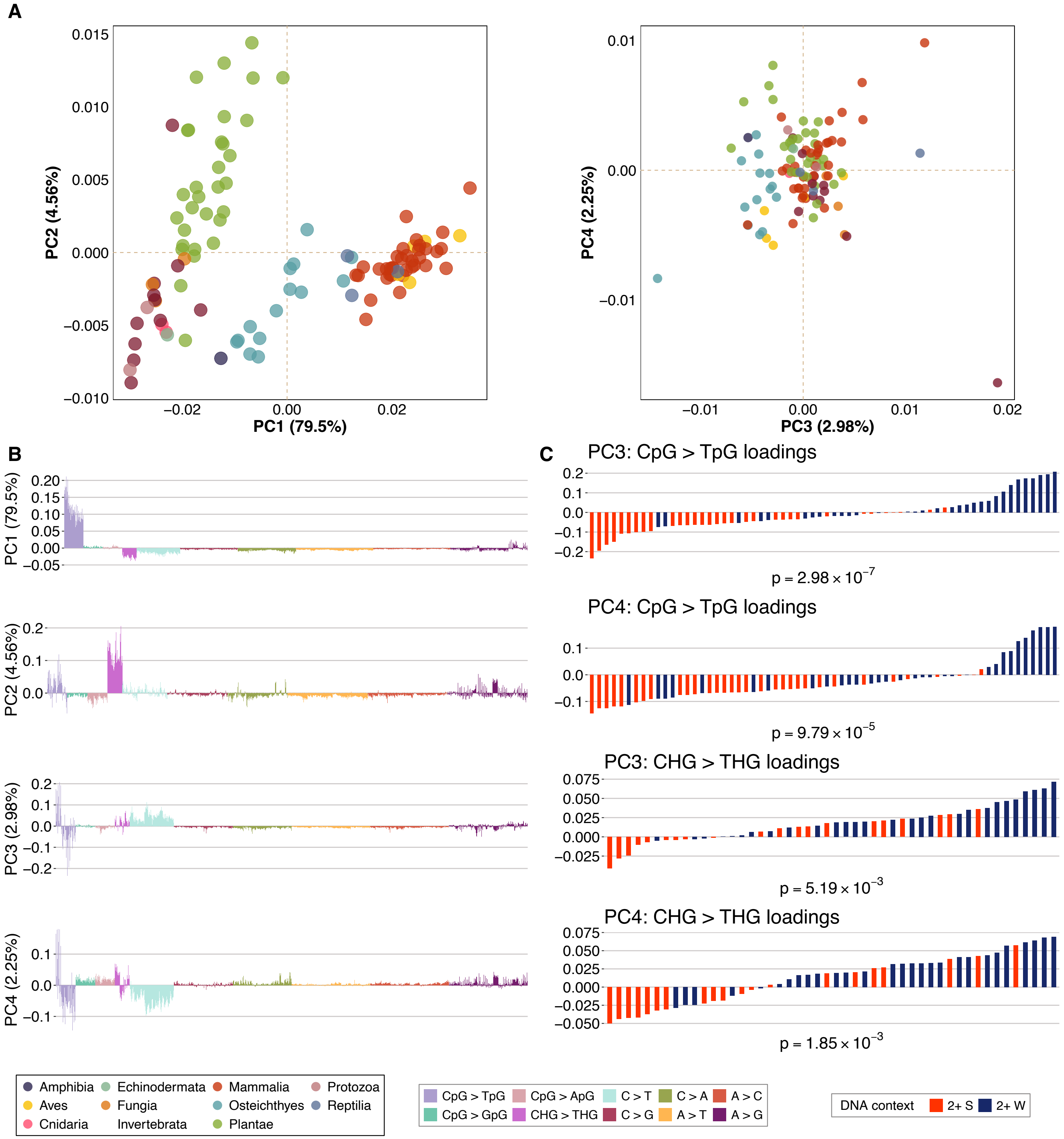


**Fig. S5** | Unscaled principal component analysis (PCA) on 5-mer fitted rates. **A**) Third and fourth principal component weights for each species shown as points, colored by assigned clade. **B**) Third and fourth principal component loadings, showing individual bars for each mutation type, colored by assigned focal mutation type labels. **C**) CpG > TpG mutation loadings for third and fourth principal components, showing individual bars for each CpG > TpG and CHG > THG mutation types, colored by surrounding nucleotide contexts (W = A/T; S = G/C). P-values correspond to Wilcoxon rank sum test results for the alternative hypothesis that loadings of CpG/CHG contexts with at least two strong nucleotides (C/G) are different than loadings of CpG/CHG contexts with at least two weak nucleotides (A/T).

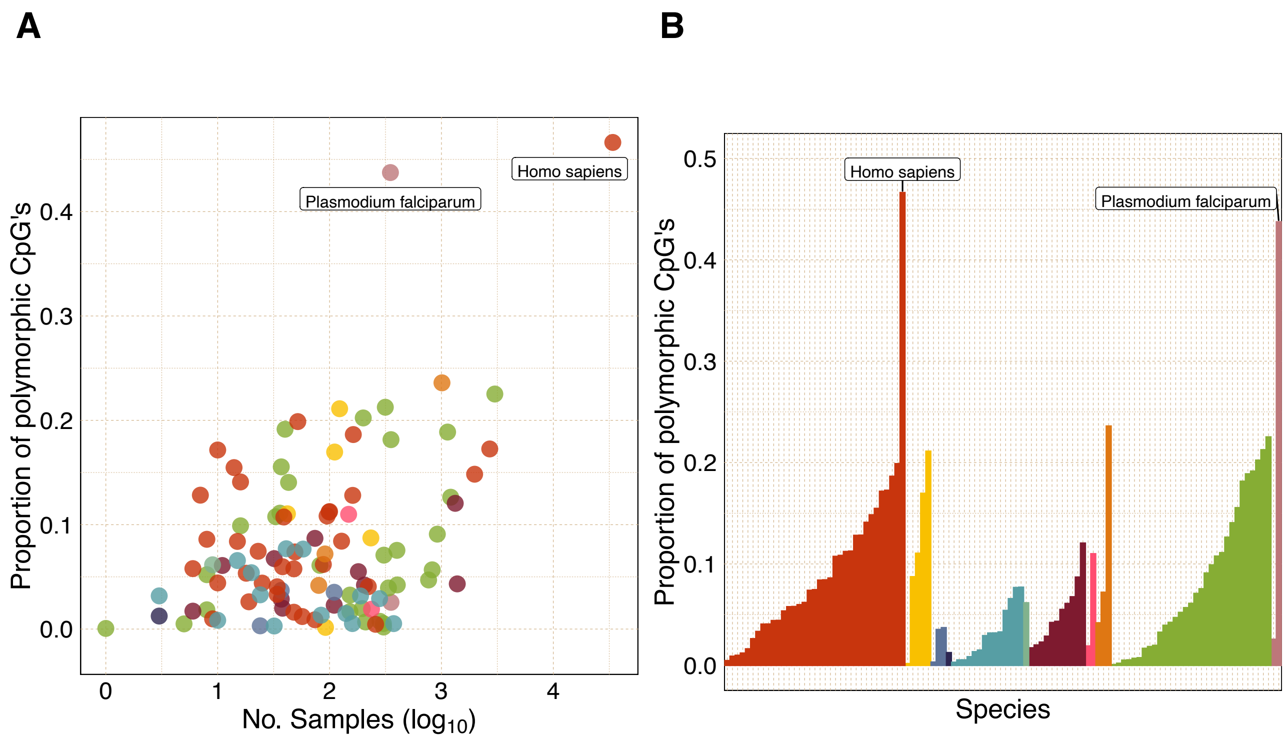


**Fig. S6** | Proportion of polymorphic CpG sites. **A)** Scatterplot showing the proportion of CpG sites that are polymorphic in non-coding regions versus the $\log_{10}$-transformed sample size of the dataset, each point shows a species colored by clade. **B)** Bar plot showing the proportion of CpG sites that are polymorphic in non-coding regions for each species. Species are arranged by clade and proportion.

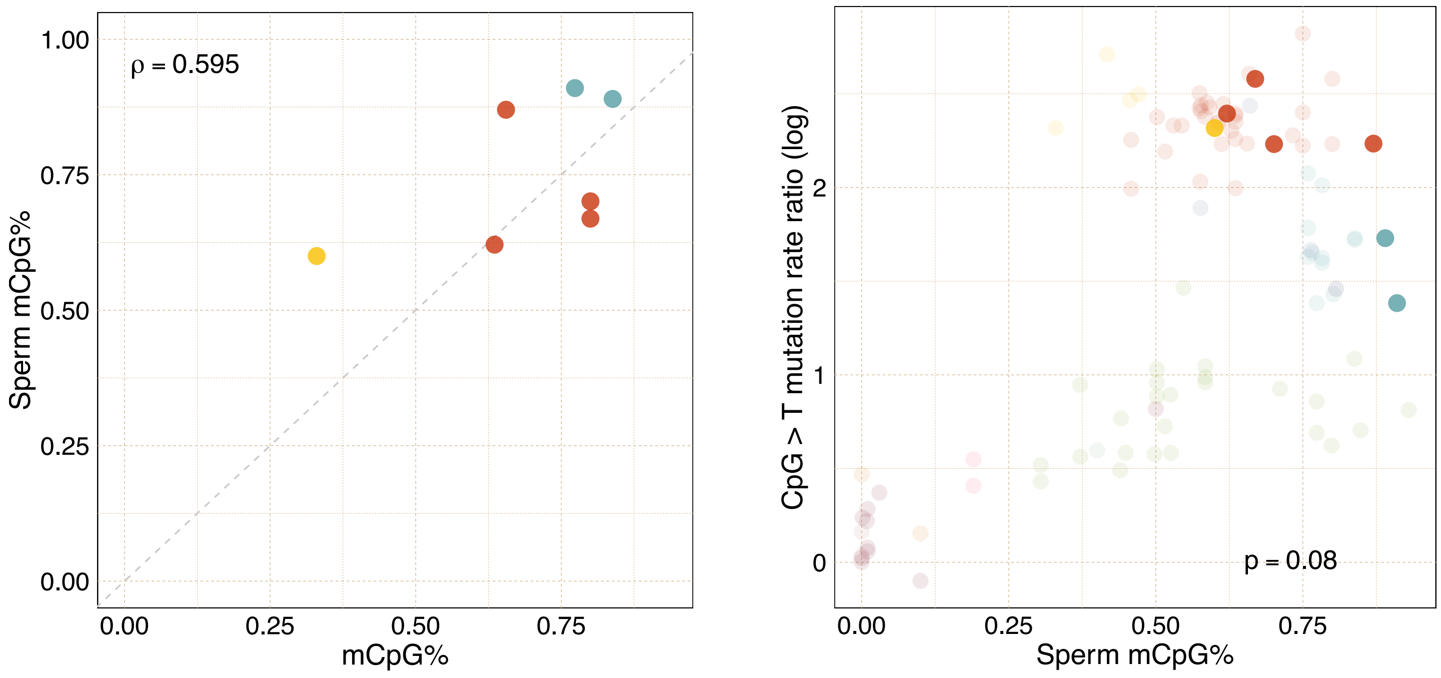


**Fig. S7** | Somatic vs sperm CpG methylation. **A)** Scatterplot of genome-wide average CpG methylation (%) in sperm tissue (y-axis) versus somatic tissues (x-axis). Spearman rho = 0.595. **B)** CpG > T mutation rate ratio vs genome-wide average CpG methylation in sperm tissue (Panel B from figure 1 is shown in high transparency). P-value corresponds to results from phylogenetic regression.

| Regression model (O/E CpG ~ CpG > T mutation rate ratio) | Number of Species | CpG > T mutation rate ratio ($\beta$) | Standard Error | Bootstrap Lower CI (95%) | Bootstrap Upper CI (95%) | p-value |
|---|---|---|---|---|---|---|
| All species | 108 | -0.125 | 0.034 | -0.197 | -0.058 | **4.03×10[-4]** |
| Vertebrates | 58 | -0.074 | 0.029 | -0.128 | -0.017 | **0.012** |
| Mammals | 35 | -0.073 | 0.038 | -0.154 | 0.002 | 0.064 |
| Bony fish | 14 | -0.097 | 0.067 | -0.220 | 0.026 | 0.168 |
| Invertebrates | 11 | -0.967 | 0.340 | -1.557 | -0.352 | **0.019** |
| Plants | 31 | -0.040 | 0.072 | -0.186 | 0.104 | 0.579 |

**Table S1** | Within-clade associations between CpG O/E ratio and CpG > T mutation rate ratio. P-value corresponds to results from phylogenetic regression. Tests were performed for clade with at least 10 species.

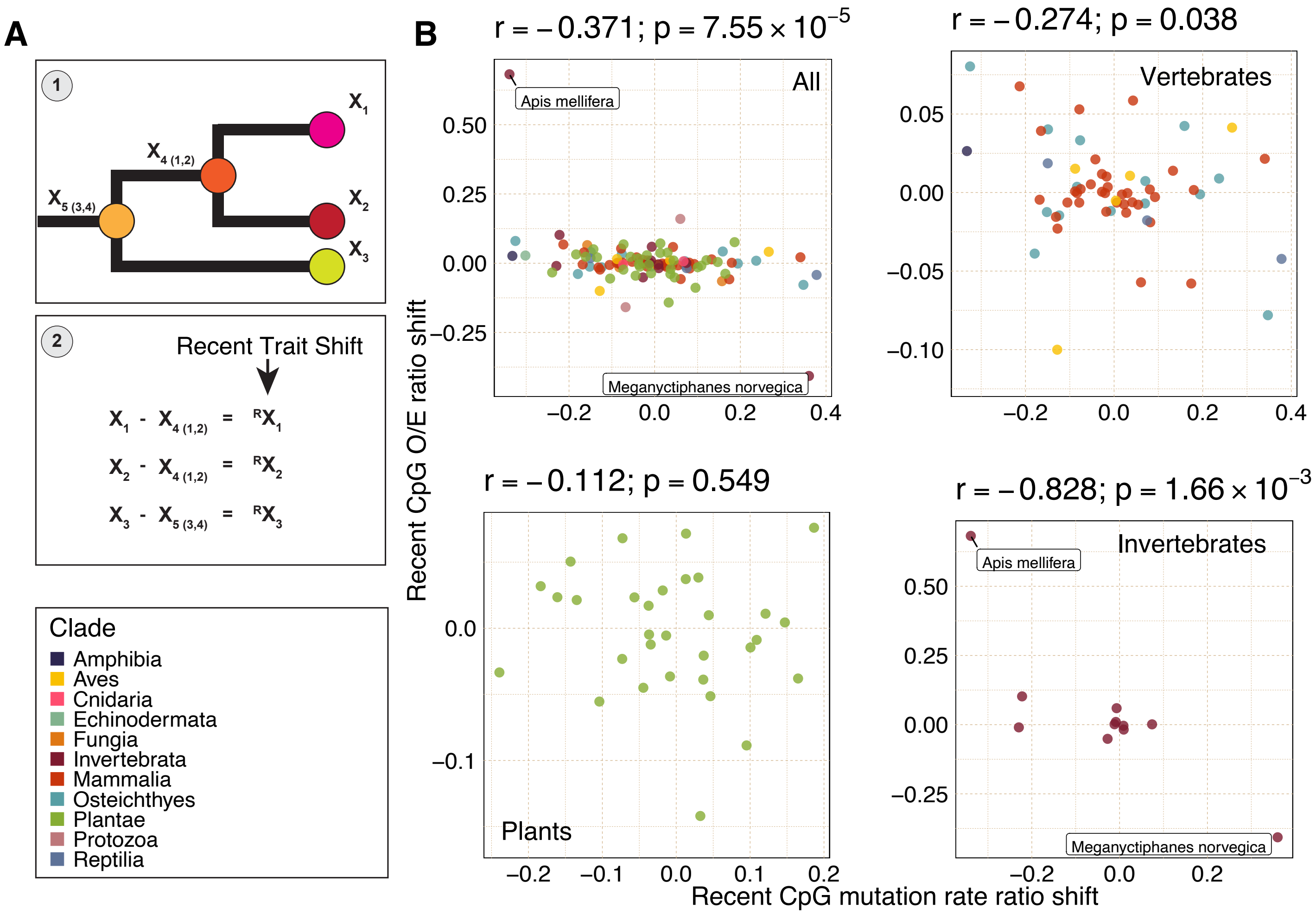


**Fig. S8** | Correlation between recent shifts in CpG O/E ratio and CpG > T mutation rate ratio. **A**) Schematic for how recent trait shifts were calculated; 1) Trait values for all internal tree nodes are estimated using fastAnc() from phytools in R4.2, 2) Recent trait shifts are defined as the difference between the extant species' trait value and its parent node's trait value. **B**) Scatterplots for recent shifts in CpG O/E ratio vs CpG mutation rate ratio. Each point shows one species, colored by assigned clade. Pearson coefficients and p-values are shown for each species grouping test (All, vertebrates, plants, and invertebrates).

| **Correlation test between recent shifts in CpG O/E ratio and CpG>T mutation rate ratio** | **Number of Species** | **Pearson's coefficient (*r*)** | **95% CI Lower** | **95% CI Upper** | **P-value** |
|---|---|---|---|---|---|
| All species | 108 | -0.371 | -0.524 | -0.196 | **$7.55\times10^{-5}$** |
| Vertebrates | 58 | -0.274 | -0.497 | -0.017 | **0.038** |
| Mammals | 35 | -0.252 | -0.540 | 0.089 | 0.144 |
| Bony fish | 14 | -0.454 | -0.794 | 0.101 | 0.103 |
| Invertebrates | 11 | -0.828 | -0.954 | -0.452 | **$1.66\times10^{-3}$** |
| Plants | 31 | -0.112 | -0.448 | 0.252 | 0.549 |

**Table S2** | Correlation tests between recent shifts in CpG O/E ratio and CpG > T mutation rate ratio. Tests were performed for clade with at least 10 species.

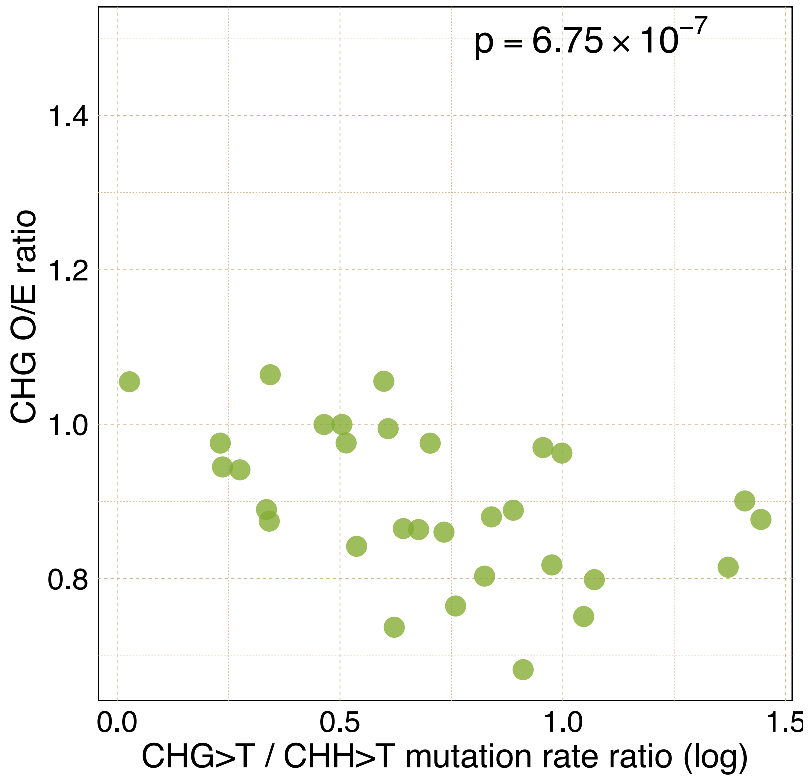


**Fig. S9** | Observed-over-expected CHG content ratio vs CHG>T mutation rate ratio in plants. Points show each plant species. P-value corresponds to phylogenetic regression of CHG O/E ratio on log-transformed CHG > T / CHH > T mutation rate ratio.

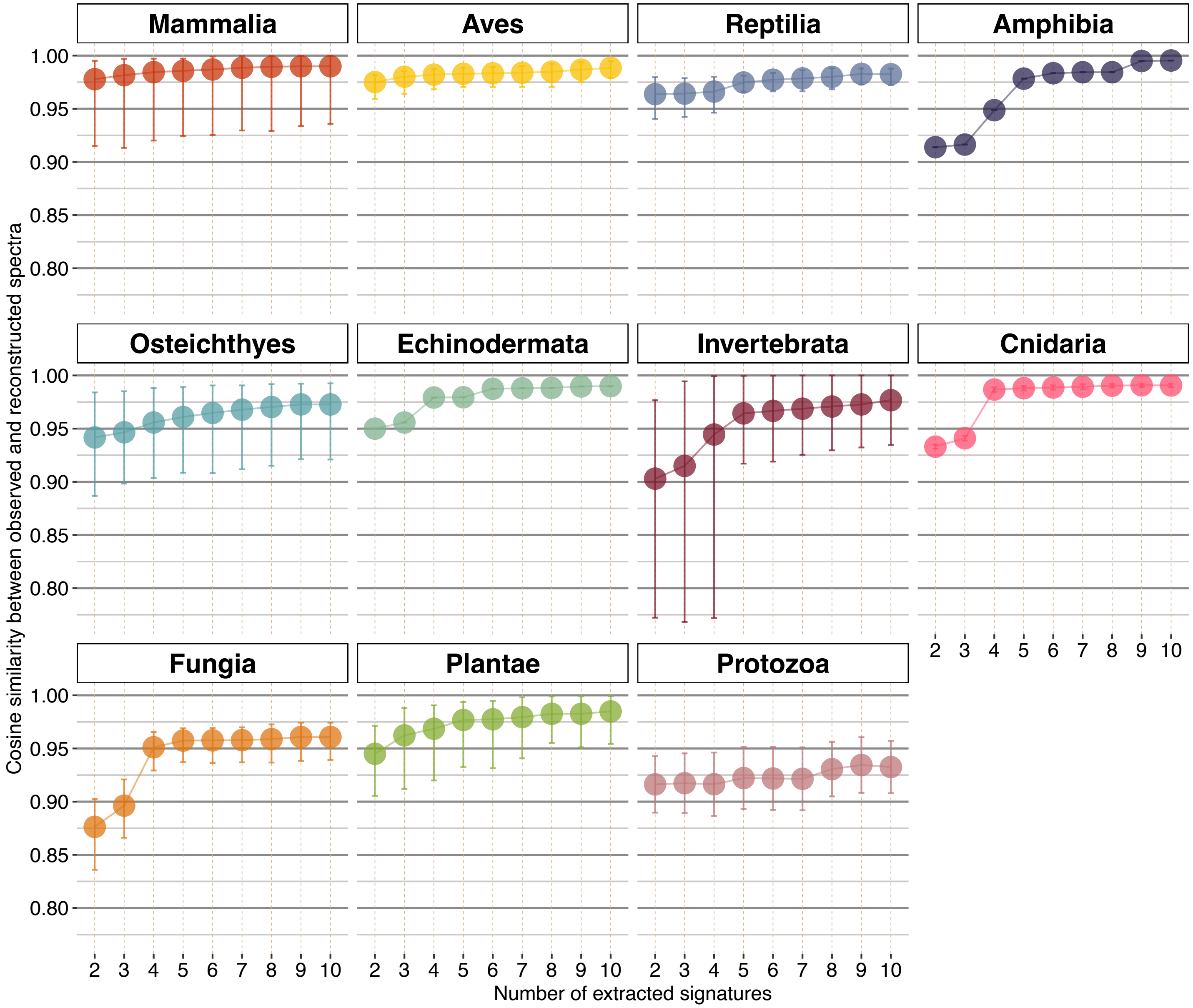


**Fig. S10** | SigFit mutational signature analysis. SigFit polymorphism spectra reconstruction performance for k extracted signatures (k=2,3,...,10). Individual points show the average cosine similarity between observed and reconstructed polymorphism spectra for each clade, with bars depicting the highest and lowest cosine similarities.

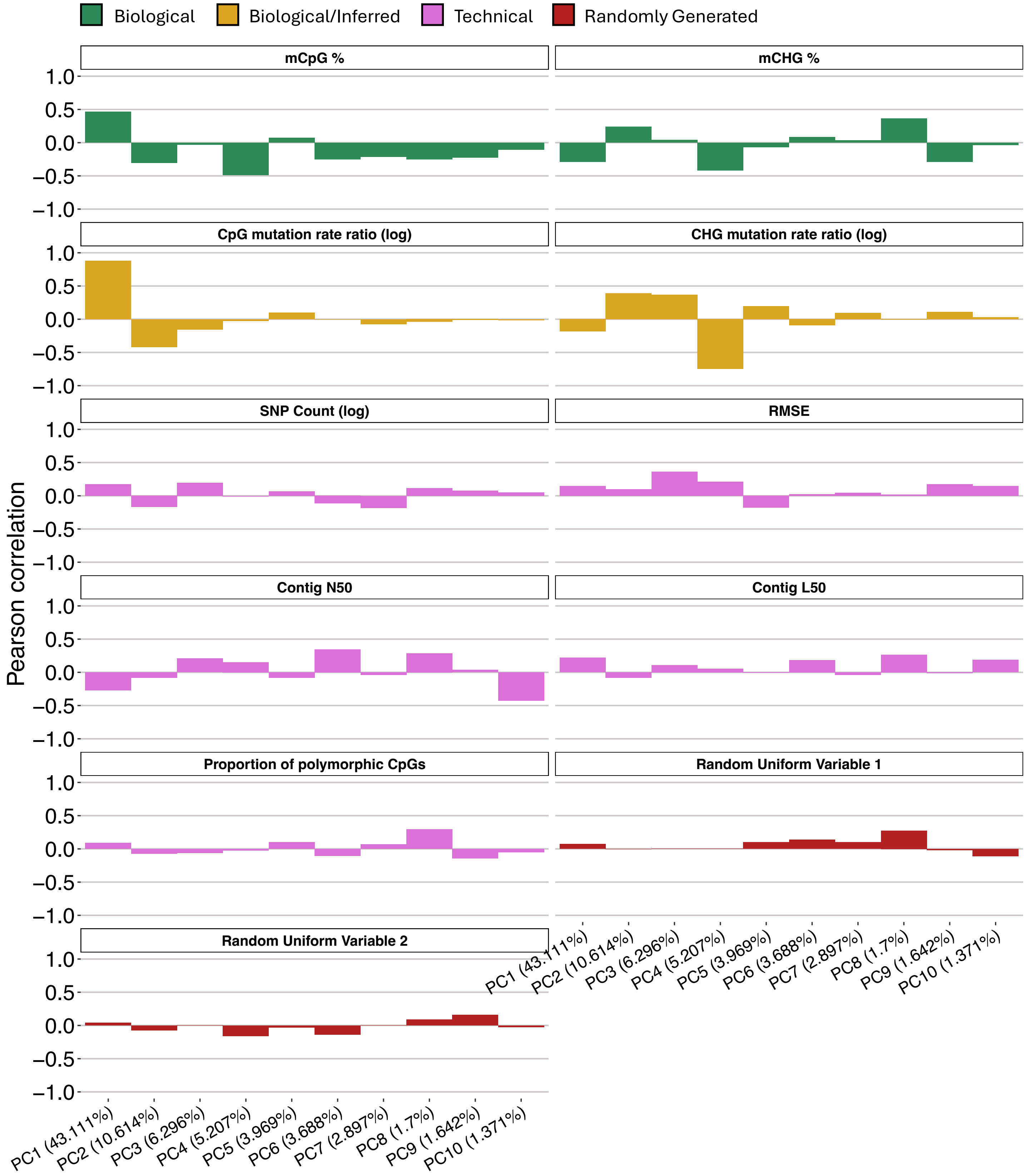


**Fig. S11** | Correlation analysis of top ten principal components from scaled 5-mer PCA against biological, technical, and generated random variables. Random variables 1 and 2 were randomly generated using the runif() function in R.
